# Phenotypic heterogeneity drives phage-bacteria coevolution in the intestinal tract

**DOI:** 10.1101/2023.11.08.566301

**Authors:** Nicolas Wenner, Anouk Bertola, Louise Larsson, Andrea Rocker, Nahimi Amare Bekele, Chris Sauerbeck, Leonardo F. Lemos Rocha, Valentin Druelle, Alexander Harms, Médéric Diard

## Abstract

Phenotypic heterogeneity in bacteria can generate reversible resistance against various stressors, including predation by phages. This allows mixed populations of phenotypically resistant and sensitive bacteria to coexist with virulent phages. However, it remains unclear if these dynamics prevent the evolution of genetic resistance in bacteria and how they affect the evolution of phages. In this work, we focus on bistable alterations of the O-antigen (known as phase variation) in *Salmonella* Typhimurium (*S*.Tm) to study how heterogeneous phenotypic resistance affects phage-bacteria coevolution. Our findings reveal that phase variation allows a stable coexistence of *S*.Tm with a virulent T5-like phage *in vitro*. This coexistence is nevertheless short-lived when *S*.Tm and the phage interact within the intestinal tract of mice. In this context, the phage evolves to also infect phenotypically resistant *S*.Tm cells, incidentally altering infectivity on other *Salmonella* serovars. In return, the broader host range of the evolved phages drives the evolution of genetic resistance in *S*.Tm, which results in phage extinction. This work demonstrates that phenotypic heterogeneity profoundly influences the antagonistic coevolution of phages and bacteria, with outcomes intricately tied to the ecological context.

## Intro

Phenotypic heterogeneity, wherein subsets of cells express distinct phenotypes from identical genomes, can have a decisive impact on the evolutionary dynamics of bacterial populations **[1]**. Phenotypically tolerant cells can for example rescue populations exposed to lethal doses of antibiotics, thereby fostering the evolution of genetic resistance **[2, 3]**. Here, we investigate the role of phenotypic heterogeneity in managing the most ancient and widespread burden on bacterial fitness: the ubiquitous, diverse and highly evolvable bacteriophages (phages). Genetic resistance and phenotypic heterogeneity can both protect bacteria against virulent phages, yet the predominant mechanism and the influence of the ecological context remain unclear. While altering or losing the phage receptor— the ligand that phages use to firmly attach to bacteria—is a common resistance strategy **[4]**, such mutations in conserved features like membrane transporters or flagella can be detrimental in the absence of phages **[5]**. Alternatively, the masking of phage receptors with surface glycans, such as capsule, O-antigen, and cell-wall glycopolymers can hinder phage adsorption with lower repercussions on the fitness of the bacteria **[6]**. The expression of glycan-modification systems is often regulated by epigenetic toggle switches, which fosters phenotypic heterogeneity **[7]**. This process, known as phase variation, generates subsets of cells phenotypically resistant to phages affected by surface glycan modifications **[8, 9]**. Every generation, cells can switch between phenotypically phage-sensitive and resistant states. This facilitates the coexistence with phages that replicate on sensitive cells—a pivotal aspect of the phage-bacteria interaction with significant implications for the design of effective phage therapy **[10–13]**. The evolutionary stability of coexistence and the broader impact of phenotypic heterogeneity on phage-bacteria coevolution nonetheless remained to be investigated.

To fill this knowledge gap, we studied the coevolution of a strain of *Salmonella* enterica serovar Typhimurium (*S*.Tm) able to modify the composition of its O-antigen by phase variation and the virulent T5-like phage ф37 **[14]**. Given that the outcome of phage-bacteria coevolution depends on the ecological context **[15]**, we compared the dynamics of phage-bacteria interactions *in vitro* and during colonization of the mouse intestinal tract. This is important because *S*.Tm is a model organism for entero-pathogens that are relevant targets for phage therapy in the intestinal tract, a promising but perfectible alternative to antibiotics requiring deeper mechanistic and evolutionary understanding **[16]**. Since many ecological factors influence the phage-bacteria interactions in the gut, the evolutionary outcomes remain difficult to predict **[17–20]**.

We found that phase variation of the O-antigen in *S*.Tm can prolong the coexistence with the phage during experimental evolution *in vitro*. However, the selection regime in the intestinal tract of mice favors the accumulation of mutations in the phage that increase its host range to all O-antigen phase variants. This drives the rapid evolution of genetic resistance in *S*.Tm and destabilizes the coexistence between phages and bacteria.

## Results

### O-antigen phase variation in S.Tm generates phenotypic resistance against ***ф***37

In non-typhoidal *Salmonella*, the outermost glycan layer is the O-antigen, a repetitive polymer of hetero-glycans attached to the lipopolysaccharide (LPS) core that hides outer-membrane proteins **[21]**. The reference strain *S*.Tm SL1344 (*S*.Tm WT) used in this study harbors two O-antigen modification systems controlled by epigenetic switches (**Figure 1A**). Firstly, the *gtrABC* operon encodes a glucosyltransferase that branches an extra glucose residue onto the galactose of the O-antigen backbone. This shifts the serotype of *S*.Tm from O:12 to O:12-2 **[22]**. Secondly, the *opvAB* operon reduces the length of the O-antigen **[23]**. Dam-dependent methylation of specific GATC sites in their respective promoters conditions the expression of these operons. The methylation patterns can change at every replication of the chromosome turning the expression ON or OFF. Importantly, silencing methylation patterns occur more frequently than patterns allowing expression, which means that most *S*.Tm WT cells do not express *gtrABC* or *opvAB* in the absence of selection for the modified O-antigen **[23, 24]**.

**Figure 1.**
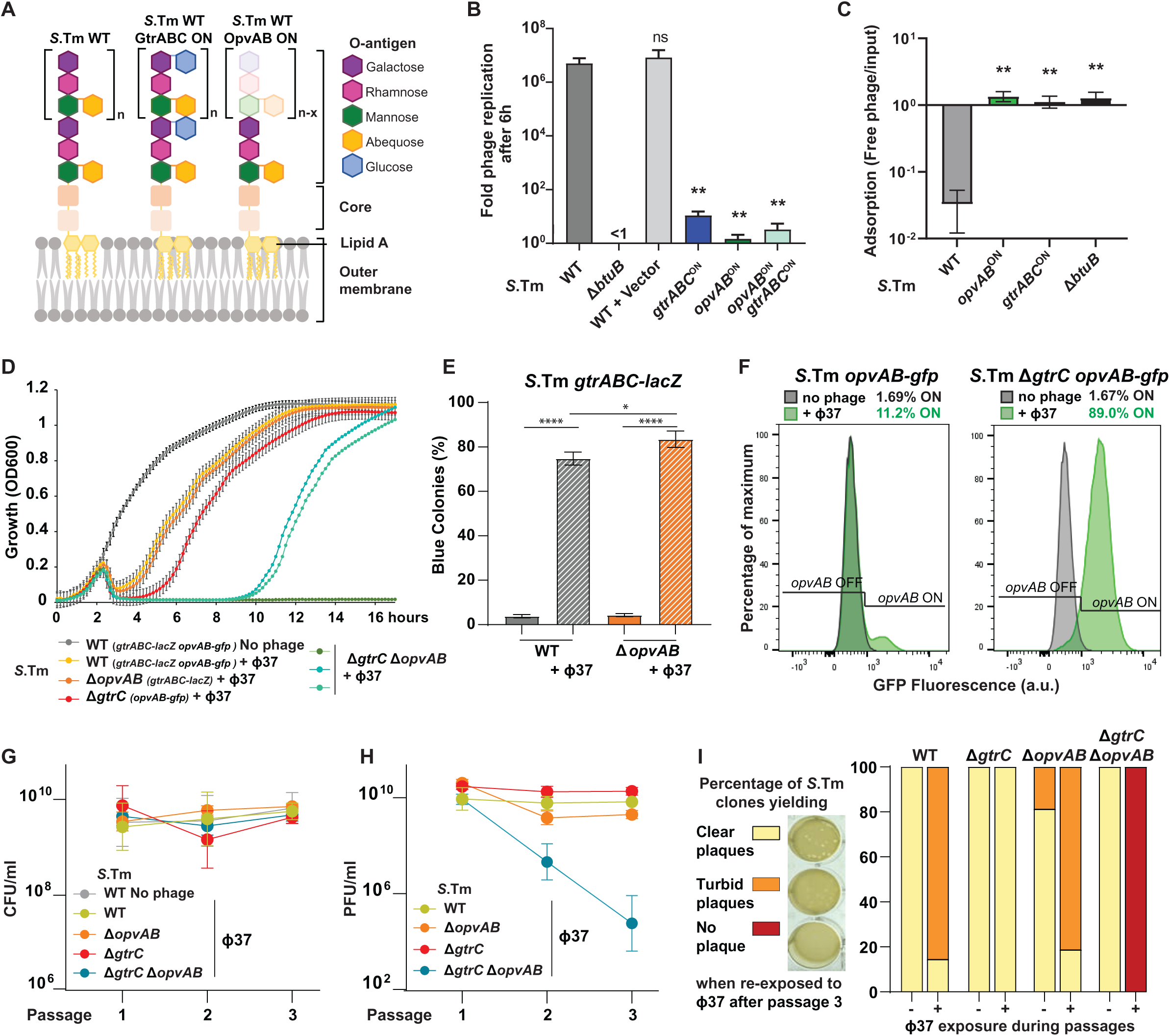
O-antigen phase variation confers phenotypic phage resistance and promotes the coexistence of *S.*Tm with ф37. **A**. O-antigen structure and modifications in *S*.Tm SL1344 (*S*.Tm WT). The GtrABC system glucosylates the O-antigen, while the OpvAB system shortens the O-antigen. **B**. ф37 replication is inhibited by O-antigen modification in *S*.Tm *gtrABC*^ON^ (carrying plasmid p*gtrABC*) and/or *opvAB*^ON^. The strain lacking the ф37 receptor BtuB (Δ*btuB*) was used as a negative control. WT+Vector corresponds to *S*.Tm WT carrying a plasmid with the same backbone as p*gtrABC*. **C**. ф37 adsorption was impaired on *S*.Tm *gtrABC*^ON^ or *opvAB*^ON^ and on the Δ*btuB* mutant. Replication and adsorption assays were performed with 6 biological replicates and each group was compared to the corresponding WT control (Mann-Whitney tests: \**p*<0.05, \*\**p*<0.01, ns: not significant). **D**. *S*.Tm WT, Δ*gtrC* and Δ*opvAB* carrying the transcriptional fusions *gtrABC-lacZ* and/or *opvAB-gfp* and the double Δ*gtrC* Δ*opvAB* mutant were inoculated in LB with or without ф37 (MOI=0.01, *n=*3) and growth kinetics were monitored for 17 hours. **E.** From the same cultures, the *gtrABC* ON/OFF status was determined by plating the bacteria on LB agar X-Gal (unpaired *t* tests: \**p*<0.05, \*\*\*\**p*<0.0001) and the *opvAB* ON/OFF status was determined by flow cytometry (**F**). **G-H**. *S*.Tm WT (*n*=9), Δ*gtrC* (*n*=6), Δ*opvAB* (*n*=6) and Δ*gtrC* Δ*opvAB* (*n*=6) were grown in the presence of ф37 (MOI=0.01) in LB. The cultures were diluted 1000-times and bacterial **(G)** and phage loads **(H)** were measured during 3 passages (see Methods). As a control, *S*.Tm WT was also passaged in the absence of ф37 (*n*=3). **I**. The susceptibility to phage ф37 was determined by plaque assay for 8 individual clones isolated from each culture. “Clear plaques” revealed susceptible clones (yellow), “no plaques” was due to mutations in *btuB* conferring full resistance (red), while “turbid plaques” reflected the expression of *gtrABC* in a large part of the population, as depicted in **Figure S1H** (orange). Results of the plaque assays are detailed in **Figure S1G**.

We first thoroughly characterized the impact of phase variation on the interaction between *S*.Tm and the T5-like virulent phage ф37 in LB medium. The phage ф37 can infect *S*.Tm cells that produces the canonical O-antigen serotype O:1,4**[5]**,12 and, like most T5-like phages, uses the membrane protein BtuB as receptor **[14]**. To assess the impact of O-antigen modifications on the replication of ф37 we used constitutive expression of *gtrABC* (*S*.Tm *gtrABC*^ON^ **[14]**) and *opvAB* (*S*.Tm *opvAB*^ON^ **[8]**) (**Table S1**). We found that the expression of each of the two phase variation systems prevented the replication of ф37 (**Figure 1B**) by limiting phage adsorption (**Figure 1C**). This was congruent with previous reports demonstrating that O-antigen glucosylation by GtrABC protects *S*.Tm against the T5-like phage SPC35 **[9]** and that O-antigen modification by OpvAB increases resistance to several phages **[8]**.

Growth dynamics showed that *S*.Tm exposed to ф37 quickly recovered after an initial decrease of the optical density (OD) (**Figure 1D**). We hypothesized that the fast recovery was due to the growth of phenotypically resistant cells after killing of the population sensitive to the phage. To confirm the selection of phenotypically resistant ON cells by the phage, we monitored the expression of the transcriptional fusions *gtrABC*-*lacZ* and *opvAB*-*gfp*. As expected, the exposure to ф37 shifted the serotype of *S*.Tm from O:12 to O:12-2 (**Figure S1A**) and increased the fraction of blue colonies on LB agar X-Gal (**Figure 1E and S1B**), meaning that *gtrABC-lacZ* was expressed. Pressure from the phage also increased the fraction of cells expressing *gfp* in *S*.Tm Δ*gtrC opvAB*-*gfp* (**Figure 1F**). Accordingly, the constitutive expressions of *gtrABC* or *opvAB* fully protected *S*.Tm against the phage (**Figure S1C**). Moreover, when both O-antigen modification systems were deleted in the double mutant Δ*gtrC* Δ*opvAB*, the regrowth of the bacteria was either not observed at all or strongly delayed (**Figure 1D**). The sequencing of isolated clones in these cases revealed the presence of *Salmonella* mutants that do not produce the wild-type BtuB **(Dataset S2**). The selection of *btuB* mutants in the absence of *gtrC* and *opvAB* demonstrated that no other mechanism generating phenotypic resistance could protect *S*.Tm against ф37 in these conditions.

### Phenotypic resistance in S.Tm allows coexistence with ***ф***37

To further investigate the impact of phenotypic resistance on the evolutionary dynamics of *S*.Tm and ф37, we repeatedly diluted batch cultures in LB medium every ten generations. Selective plating and plaque assays determined bacterial counts and phage titers respectively (**Figures 1G and H**). Phenotypic resistance in *S*.Tm WT, and in the mutants Δ*gtrC* and Δ*opvAB* allowed phages and bacteria to coexist for at least 30 generations (i.e., three passages) despite bottlenecks generated by repeated dilutions of the cultures. In the double mutant Δ*gtrC* Δ*opvAB*, the impossibility to escape ф37 by O-antigen phase variation led to the emergence of genetically resistant *btuB* mutants (**Dataset S2**). In these cases, we observed a sharp decrease of the phage population unable to overcome the dilution bottlenecks (**Figure 1H**). A similar trend was observed when phenotypic resistance was made constitutive, which drastically limits phage replication (**Figure S1D**). This is in accordance with the theoretical prediction that the ON to OFF switch rate must be high enough to generate a susceptible population of bacteria able to sustain phage replication as in *S*.Tm WT **[13]**. Longer experiments comprising 20 passages showed that the phage-bacteria coexistence mediated by phase variation could last for at least 200 bacterial generations (**Figure S1E and F**).

To confirm that phase variation prevents the selection of *btuB* mutants during phage exposure, we assessed the resistance to ф37 in isolated clones of *S*.Tm after 3 passages by plaque assay (**Figures 1I and S1G**). We analyzed eight clones from two lines without phage and six lines with phage (**Figure S1G**). ф37 produced turbid plaques when the lawns were generated from cells able to express *gtrABC* and pre-exposed to the phage. This is because these lawns contained mixtures of phenotypically resistant *gtrABC* ON cells and sensitive OFF cells **[14]**, as confirmed by the analysis of ф37 plaques on clones picked according to the expression of *gtrABC*-*lacZ* revealed on LB agar X-Gal (**Figure S1H**). No visible plaques suggested genetic resistance, and amplicon sequencing confirmed mutations in *btuB* (**Dataset S2**). Clear plaques were generated from clones sensitive to ф37. No resistant clones were detected from the single mutant *S*.Tm Δ*gtrC* because OpvAB protects *S*.Tm from the phage (**Figure 1B**). However, in this case, no turbid plaques were observed. This can be explained by the high ON to OFF switching frequency of *opvAB* expression **[23]**: during growth in the absence of phages before the plaque assay, newly formed *opvAB* OFF cells quickly replace the *opvAB* ON cells, hence generating fully sensitive bacterial lawns on which ф37 makes clear plaques. The fixation of resistant *btuB* mutants in *S*.Tm Δ*gtrC* Δ*opvAB* correlated with the sharp decrease of the phage titer (**Figure 1H**), demonstrating that phage-bacteria coexistence was not possible without phase variation.

### Phenotypic resistance delays the fixation of btuB mutants when S.Tm is exposed to ***ф***37 during intestinal colonization

Next, we investigated the impact of phase variation in the intestinal tract where ecological heterogeneity already favors phage-bacteria coexistence to some extent, by limiting the physical interaction between bacteria and phages **[20]**.

To study the phage-bacteria coevolution in the gut, we performed long-term infections in mice (**Figure 2A**). For this, we used the attenuated strain *S*.Tm Δ*ssaV* (S.Tm* **[25]**) and mice harboring a simplified intestinal microbiota that cannot exclude *Salmonella* **[26]**. These conditions allow stable intestinal colonization by *S*.Tm* for 10 days **[27]**. Bacteria and phages were detected in homogenized fecal samples respectively by selective plating (**Figure 2B**) and plaque assay (**Figure 2C**). We compared the outcomes of phage-bacteria coevolution using phase variable *S*.Tm* and the double mutant *S*.Tm* Δ*gtrC* Δ*opvAB*.

**Figure 2.**
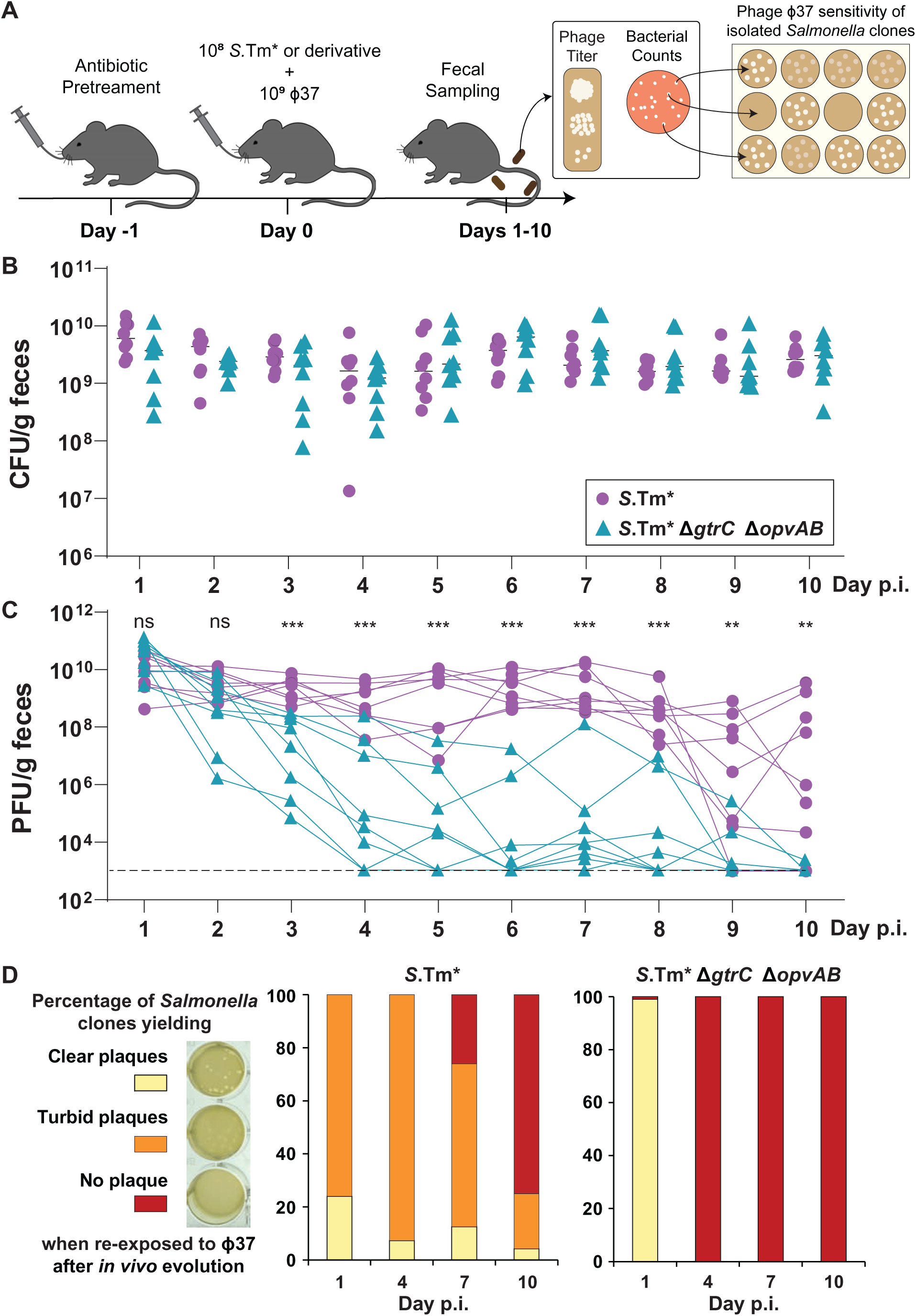
The phage-bacteria coexistence is short-lived during intestinal infection. **A.** Experimental setup. Streptomycin pretreated C57BL/6 Low Complexity Microbiota (LCM) mice were infected with the attenuated *S*.Tm Δ*ssaV* strain (*S.*Tm*, 10^8^ CFUs, 8 mice) or with the *S.*Tm* derivative mutant Δ*gtrC* Δ*opvAB* (8 mice). Mice from both groups received 10^9^ PFU of ф37 30 min after *S*.Tm inoculation. Bacterial **(B)** and phage loads **(C)** were quantified in fecal samples for 10 days (Mann Whitney tests: \**p*<0.05, \*\**p*<0.01, \*\*\**p*<0.001. ns: not significant). **D.** Phage susceptibility of 12 clones isolated from each mouse at day 1, 4, 7 and 10 post-infection (p.i.) was determined by plaque assay. Yellow = clear plaques, susceptible clone; Orange = turbid plaques, heterogeneous phenotypic resistance linked to the expression of *gtrABC*; Red = no plaque, resistance linked to mutations in *btuB*. Results of the plaque assays are detailed in **Figure S2**.

We first observed that the phage did not prevent the colonization of the gut by *S*.Tm*. The intestinal *Salmonella* loads were comparable with and without phase variation (**Figure 2B**). On the other hand, the phage titers decreased rapidly in the absence of phenotypic resistance when *S*.Tm* Δ*gtrC* Δ*opvAB* was the host of ф37. We performed plaque assays to test the resistance of isolated *Salmonella* clones after 1, 4, 7 and 10 days following infection. Twelve clones from eight mice per condition were analyzed (**Figures 2D and S2**). The fixation of resistant *btuB* mutants in *S*.Tm* Δ*gtrC* Δ*opvAB* correlated with the extinction of the phage (**Figure 2D**, **right panel**), as observed *in vitro* (**Figure 1I**). Intriguingly, although the phage-bacteria coexistence was more stable with S.Tm*, the phage titers eventually diminished in half of the mice after 8 days. This was also concomitant with the rise of resistant *btuB* mutants (**Figure 2D**, **left panel**).

We verified that O-antigen phase variation in fully virulent *S*.Tm derivatives also delayed the fixation of *btuB* mutants under phage pressure, including in Δ*gtrC* and Δ*opvAB* single mutants (**Figure S3**). For this, we performed short-term infections in conventional C57BL/6 mice pretreated with streptomycin to allow robust intestinal colonization by *S*.Tm **[28]**. Exhausted LB was used as mock treatment in control groups without phage (**Figure S3A**). The bacterial and phage loads were determined in homogenized fecal samples for three days (**Figures S3B and C**). Twenty *Salmonella* clones per mouse were analyzed after 3 days post-infection. Turbid plaques formed on clones from the control group without phage showed that the expression of *gtrABC* might be slightly advantageous in the gut in the absence of ф37 (**Figure S3D**). However, *gtrABC* was expressed at higher frequencies in clones exposed to the phage (**Figures S3D and F**). No resistant mutants (i.e., clones generating no plaque) were observed in control groups, whereas in phage-treated mice, phenotypic resistance via expression of *gtrABC* or *opvAB* prevented the fixation of resistant mutants in most mice (**Figures S3D, E and F**), otherwise overrepresented in mice infected by the double mutant *S*.Tm Δ*gtrC* Δ*opvAB* (**Figure S3G**).

We concluded that phenotypic resistance mediated by phase variation plays an important evolutionary role in the intestinal tract when *S*.Tm is exposed to phages: it protects *Salmonella* from quickly losing the conserved vitamin B12 transporter BtuB, receptor of ф37. However, long-term infections also showed that the phage-bacteria coexistence was only short lived as *btuB* mutants eventually emerged in phase-variable populations. The next step was to understand this phenomenon.

### Accumulation of mutations in the lateral tail fiber protein increases the host-range of ***ф***37

Different plaque morphologies were observed with phage isolates from mice infected with *S*.Tm*, suggesting the presence of ф37 variants (**Figure 3A**). Sequencing indeed revealed non-synonymous mutations in these phages compared to the ancestor (as detailed in **Dataset S2**). Mutations were particularly frequent within the *ltf* gene, encoding the lateral tail fiber protein (**Figure 3B**). Except for a short deletion (mutation ΔA56-A97), all the non-synonymous mutations in *ltf* were found between residues 468 and 556. To further characterize the accumulation of mutations in *ltf*, we therefore sequenced amplicons of the *ltf* locus between residues 463 and 570. We randomly picked a set of 10 phages from each of the 16 mice presented in **Figure 2** on the last day phages were detected. Strikingly, all the phages in mice infected with *S*.Tm* had a mutated *ltf* allele. By contrast, in mice infected with the Δ*gtrC* Δ*opvAB* mutant (no O-antigen phase variation), only three mutations in *ltf* residues 469 and 556 were identified and 27.5% (22/80) of the phages harbored the ancestor locus.

**Figure 3.**
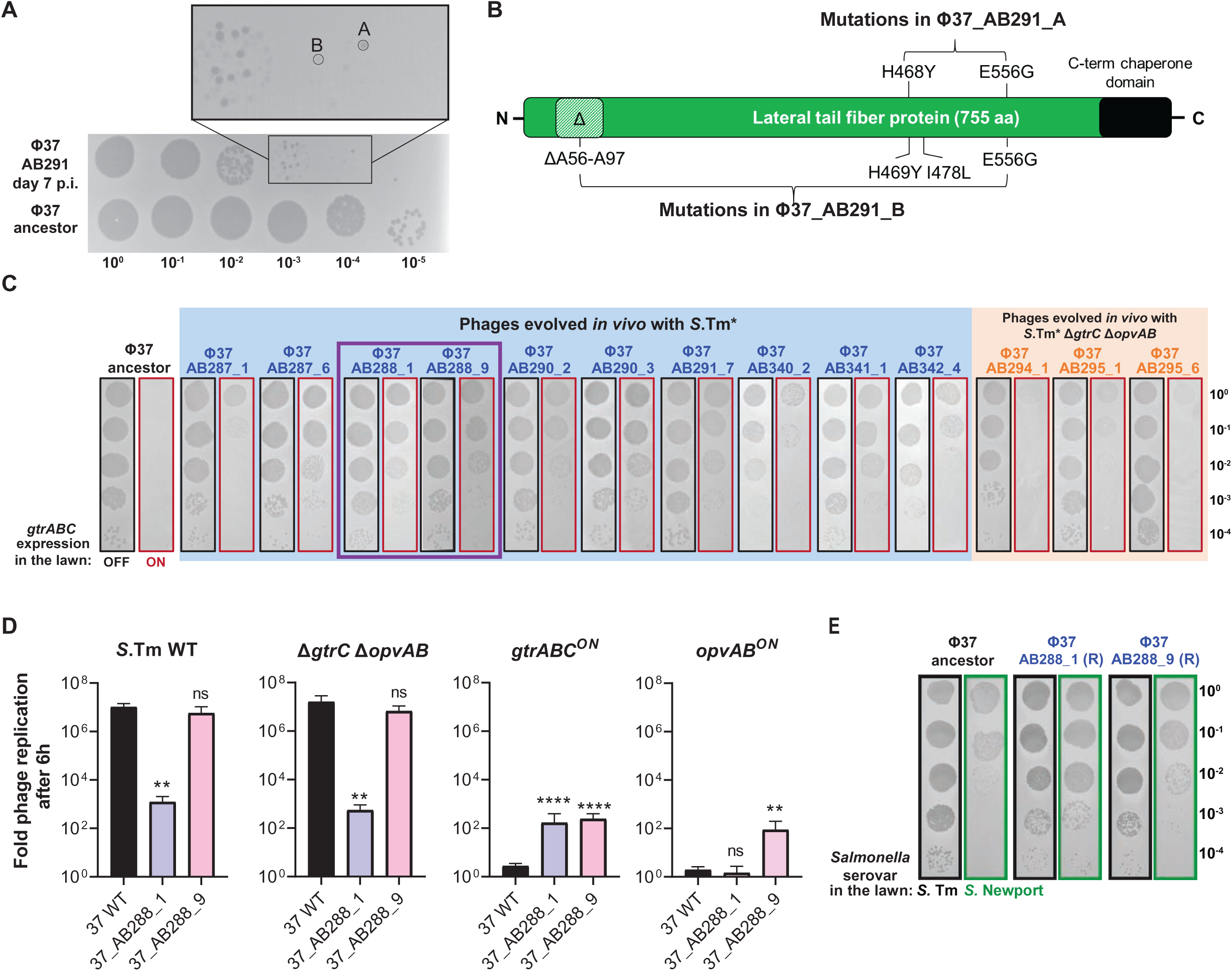
Mutations in *ltf* broaden the host range in evolved phages. ф37 accumulated mutations in the lateral tail fiber protein (Ltf) during replication in mice infected with *S*.Tm*. **A.** Phages from fecal samples were enumerated by plaque assay, revealing heterogeneous plaque morphologies. Representative plaques (mouse #AB291) were compared to plaques generated by the ancestor ф37 on a lawn of *S*.Tm WT. **B.** Two phages (ф37_AB291_A and ф37_AB291_B) that displayed different plaque morphologies were sequenced, revealing different sets of mutations in Ltf. Amino acid substitutions are indicated and “Δ” denotes a deletion. **C**. Evolved ф37 displayed an increased infectivity on *S*.Tm *gtrABC*^ON^ in comparison to the ancestor ф37. Phages isolated from mice infected with *S*.Tm* or *S*.Tm* Δ*gtrC* Δ*opvAB* (from Figure 2C) were sequenced and infectivity was tested by plaque assay on lawns of *S*.Tm WT (mainly OFF) and *S*.Tm *gtrABC*^ON^ (Δ*gtrC* Δ*opvAB* p*gtrABC*). The Ltf mutations of each phage are presented in supplementary **Figure S4A** and their full genotype is presented in **dataset S2**. **D**. *In vivo* evolved phages replicated better on *S*.Tm *gtrABC*^ON^ and *S*.Tm *opvAB*^ON^ than the ancestor ф37. The replication of two evolved phages (ф37_AB288_1 & ф37_AB288_9, framed in **C**) was tested in LB on *S*.Tm WT (*n*=6), Δ*gtrC* Δ*opvAB* (*n*=6), Δ*gtrC* Δ*opvAB* p*gtrABC* (*gtrABC*^ON^) (*n*=9) and Δ*gtrC opvAB*^ON^ (*n*=6). For each experiment, replication of the evolved phages was compared to the ancestor ф37 (Mann-Whitney tests: \**p*<0.05, \*\**p*<0.01,*** *p*<0.001, **** *p*<0.0001, ns: not significant). **E**. Ltf mutations of phage ф37_AB288_1 increased infectivity on *Salmonella enterica* serovar Newport (*S*. Newport). Ltf mutations present in phage ф37_AB288_1 and ф37_AB288_9 were transferred into the ancestor ф37 genetic background (**Figure S5**). The resulting mutants were spotted on lawns of *S*.Tm and *S*. Newport. Plaque assays with all the evolved and re-constructed ф37 on different *Salmonella* serovars are presented in **Figure S6**.

The whole genome sequencing of a subset of evolved phages confirmed the accumulation of mutations in *ltf* and revealed additional mutations in the rest of the phage chromosome (**Dataset S2**). Taken together, we have identified 10 altered *ltf* alleles in phages from mice infected with *S.*Tm* and only 3 variants in mice infected with *S*.Tm*Δ*gtrABC* Δ*opvAB* (**Figure S4A**). Some variants coexisted in the same mouse (*e.g.* evolved phages ф37-AB228_1 and AB228_9).

The lateral tail fiber is crucial in T5-like phages for specific host recognition via the reversible binding to O-antigen glycans **[29]**. We therefore hypothesized that ф37 accumulates mutations in *ltf* to overcome phenotypic resistance from phase variation of the O-antigen. Compared to the ancestor ф37, phages evolved on *S*.Tm* *in vivo* were indeed better at infecting *S*.Tm *gtrABC*^ON^ (**Figure 3C and D**) and/or *S*.Tm Δ*gtrC opvAB*^ON^ (**Figure 3D and S4B**). In certain cases, the trade-off was a reduced ability to infect the *S*.Tm OFF cells (*e.g.* in evolved ф37-AB228_1).

To demonstrate that the increased infectivity was caused by mutations in *ltf* only, we have re-constructed a subset of *ltf* mutations in the ancestor ф37 background (**Figures S5A and B**) **[30]**. The infectivity of all re-constructed *ltf* mutants was improved on *gtrABC*^ON^ bacteria compared to the ancestor phage (**Figure S5C**). Certain mutants were slightly less efficient at infecting *S*.Tm Δ*gtrC opvAB*^ON^ than the ancestor, suggesting another trade-off that could explain the coexistence of different evolved phages in some mice.

Given that GtrABC generates a substantial modification of the O-antigen of *S*.Tm that leads to serotype shift, we hypothesized that the accumulation of mutations in *ltf* could alter the host range of ф37 not only regarding *S*.Tm phase-variants but also other *Salmonella* serovars. To test this, we measured the infectivity of ф37 and the *ltf* mutants on various *Salmonella* serovars including Enteritidis and Gallinarum (both O:9), Limete (a Typhimurium-like O:4), Senftenberg (O:1,3,19), Choleraesuis (O:7), Anatum (O:3,10) and Newport (O:8) **[31]**. Interestingly, some evolved phages were better at infecting *S.* Newport (**Figure 3E and S6**). However, these alleles drastically reduced the ability to infect *S*. Gallinarum, although not Enteritidis, both O9 (**Figure S6**). All evolved phages also lost their ability to infect *S*. Senftenberg compared to ancestor. *S.* Anatum and *S.* Choleraesuis were fully resistant to all the tested phages.

In summary, these findings illustrate that ф37 has the capacity to adapt to O-antigen modifications by altering the lateral tail fiber protein through mutations. This alteration of the host range extends beyond infecting different phase variants of *S*.Tm and implies trade-offs. For instance, it can enhance the ability to infect alternative hosts like *S*. Newport while simultaneously reducing infectivity on other hosts like *S*. Gallinarum and Senftenberg.

### Evolved phages with broad host-range drive rapid evolution of *btuB* mutants during infection

Lastly, we asked if the ability of evolved phages to kill phase variants accelerates the fixation of resistant *S*.Tm *btuB* mutants (**Figure 4**). We compared the outcome of *S*.Tm* evolution in mice with either the ancestor ф37 or a mixture of evolved phages isolated from experiments described in **Figure 2** and characterized in **Figure 3**. We mixed two evolved phages that were found in a mouse infected by *S*.Tm* (Evolved phages ф37-AB228_1 and AB228_9). The rationale behind this was that evolved phages could coexist due to their distinct abilities to infect the different *S*.Tm phase variants, together consequently promoting the fixation of *btuB* mutants. Like the ancestor, the evolved phages did not reduce the population size of *S*.Tm* in the gut (**Figure 4B**). However, the intestinal loads of the evolved phages were decreasing faster than the ancestor (**Figure 4C**). This correlated with the rapid appearance of resistant mutants in the *S*.Tm* population exposed to evolved phages (**Figure 4D and S7**), demonstrating that phages able to kill *S*.Tm despite phase variation favor genetically resistant mutants, which, in turn, impairs the replication of the phages in the infected mice.

**Figure 4.**
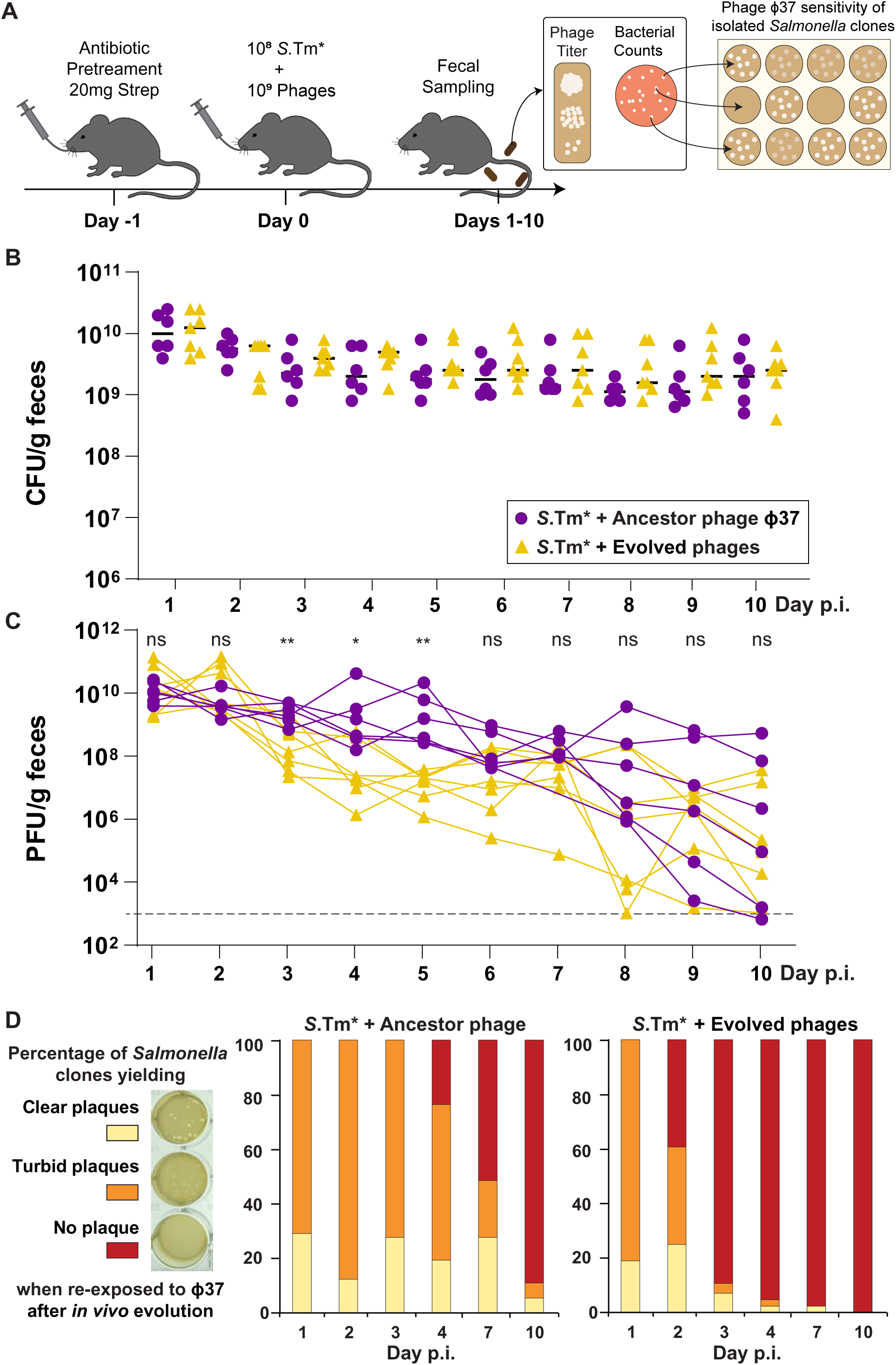
*In vivo* evolved phages accelerate the fixation of phage-resistant *btuB* mutants. **A.** Experimental setup. Streptomycin pretreated C57BL/6 LCM mice were infected with the attenuated strain *S*.Tm Δ*ssaV* (*S.*Tm*) (13 mice infected with 10^8^ CFUs). Mice received either the ancestor ф37 (6 mice) or a 1:1 mixture of the evolved phages ф37-AB288-1 + ф37-AB288-9 (7 mice), 30 min after *S*.Tm inoculation (10^9^ PFU). Bacterial **(B)** and phage loads (**C)** were quantified in fecal samples for 10 days (Mann Whitney tests: \**p*<0.05, \*\**p*<0.01, \*\*\**p*<0.001. ns: not significant). **D.** Phage susceptibility was determined by plaque assays on 12 clones isolated from each mouse at day 1, 2, 3, 4, 7 and 10 post-infection (p.i.). Yellow = clear plaques, susceptible clone; Orange = turbid plaques, heterogeneous phenotypic resistance linked to the expression of *gtrABC*; Red = no plaque, resistance linked to mutations in *btuB*. Detailed results of the plaque assays are presented in **Figure S7**.

## Discussion

In the intestinal tract, the serotype shift due to phase variation of the O-antigen enables *S*.Tm to escape neutralization by the adaptive immune system and promotes long-term colonization of the host **[14, 22]**. Nevertheless, when a polyvalent vaccine renders phase variation futile, *S*.Tm further adapts by producing a very short O-antigen at the cost of reducing its virulence and its resistance to environmental stress **[14]**, hence demonstrating that phase variation can prevent the evolution of detrimental mutations in *Salmonella*. Modifications of the O-antigen originally interfere with phage infection **[8, 9]**, therefore we reasoned that phase variation should affect the phage-bacteria interaction in the gut and preserve the bacteria from evolving potentially costly resistance, the same way it protects *S*.Tm from short O-antigen evolution under selective pressure from the immune system. The results presented in this study confirm this intuition: phenotypic resistance mediated by phase variation of the O-antigen has a profound impact on the antagonistic phage-bacteria coevolution in the intestinal tract. Along those lines, virulent phages select for phase variation in glycans produced by the intestinal bacteria *Bacteroides thetaiotaomicron* **[32]** and *Bacteroides intestinalis*, which favors the persistence of virulent phages like crAss001 in the gut **[33]**.

Nevertheless, genetic resistance eventually emerged in *S*.Tm exposed to ф37 in the gut. This occurred rapidly in the absence of protective phase variation in the double mutant Δ*gtrC* Δ*opvAB*, or once phages evolved to target the phase variants in *S*.Tm WT. Resistant mutants became predominant in as little as three days of within-host growth (**Figures S3 and 4**), highlighting the strong selective pressure exerted by phages on *S*.Tm in the intestinal environment. The mutations that resulted in resistance altered BtuB, with minimal redundancy observed between independent experiments (**Dataset S2**), hence no mutational hot spot was necessary for such fast evolution. While the phage did not completely eliminate *S*.Tm from the gut, whether phase variation was present or not, our findings imply that phages could be highly effective in counter-selecting functions that promote the growth or virulence of pathogenic bacteria in this environment. This would apply the concept of phage steering **[34]** to pathogens in the gut.

Besides evolution of *S*.Tm, the most striking finding was the evolutionary path of ф37 in the intestine. The vast majority of *in vivo* evolved phages harbored mutations in the lateral tail fiber protein that interacts with the O-antigen and determines the host range **[35]**. Mutations that change residues 469 and 556, detected in most evolved phages (**Dataset S2**), also emerged in mice infected with the double mutant *S*.Tm Δ*gtrABC* Δ*opvAB*. These mutations provide an advantage independently from phenotypic resistance presumably by increasing infectivity on cells producing the unmodified O-antigen. However, when evolving in the presence of *S*.Tm WT, the phage accumulated mutations in the lateral tail fiber protein that further improved its ability to kill the phase variants.

Restricting the host-range to the unmodified *S*.Tm cells is a prudent exploitation of the bacteria that ensures a stable production of phages, as observed for 200 bacterial generations in LB medium (**Figure S1F**) **[36]**. By killing the phase variants, “greedy” evolved phages selected for resistant *btuB* mutants, which resulted in the extinction of the phage population (**Figures 2**). Nevertheless, evolved ф37 can also more efficiently kill *Salmonella* serovar Newport (**Fig. S6**). Evolved phages could therefore thrive in more complex microbial communities in which multiple strains of *Salmonella* or related species like *Escherichia coli* could be alternative hosts. In any case, the swift adaptation of phages to phase variation observed in this study implies that phages can be “trained” via experimental evolution to bypass phenotypic resistance **[37]**. However, the choice of the environment, such as the large intestine of mice as opposed to *in vitro* passages in LB medium, could significantly enhance the effectiveness of this approach.

**Figure S1.**
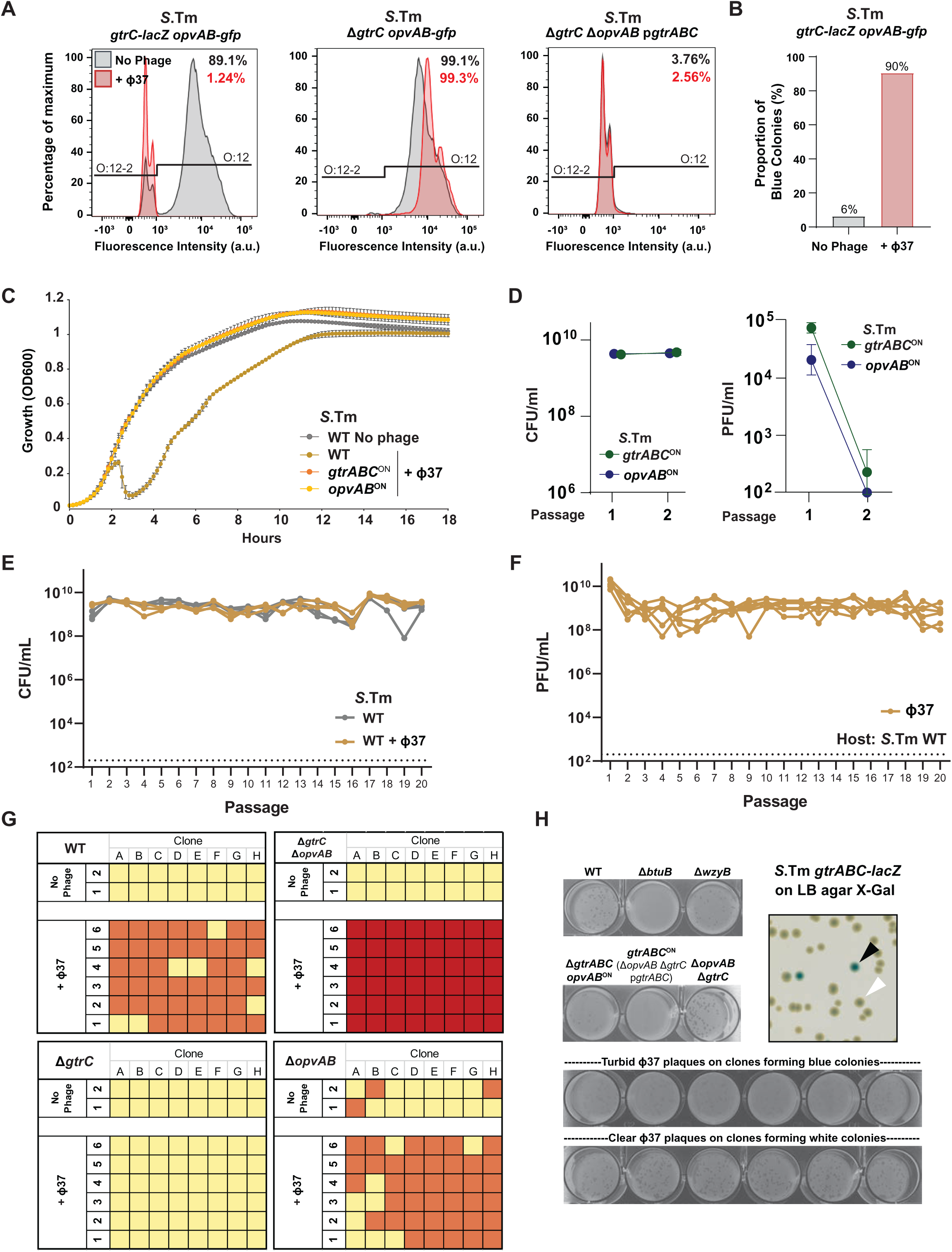
*In vitro* characterization of phage-bacteria interactions. **A.** Indirect detection of the O-antigen glucosylation by GtrABC *via* immunostaining and flow cytometry. The *S*.Tm WT and Δ*gtrC* strains (carrying the *gtrABC*-*lacZ* and/or the *opvAB*-*gfp* transcriptional fusion) were grown in the presence or absence of ф37. Bacteria were treated with the STA5 Anti-O:12 antibody that binds specifically to the unmodified *S*.Tm O-antigen. After immunostaining with the ALEXA Fluor-647-labelled secondary antibody, bacteria were analyzed by flow cytometry, revealing the proportion of cells producing the O:12 (stained) or the O:12-2 (glycosylated by GtrABC, unstained) O-antigen. The Δ*gtrC* Δ*opvAB* p*gtrABC* strain (*gtrABC*^ON^) was treated similarly and was used as a fully glycosylated O:12-2 control. **B.** *S*.Tm WT (carrying the *gtrABC*-*lacZ* transcriptional fusion) cultures from the previous experiment were plated on LB agar X-Gal, revealing the proportion of *gtrABC* ON cells after being exposed to phage ф37. **C**. Growth kinetics of *S*.Tm and derivative strains with and without ф37 (MOI=0.01, *n*=3) were monitored over 18h. Cells constitutively expressing *gtrABC* (*gtrABC*^ON^) or *opvAB* (*opvAB*^ON^) were fully resistant to ф37. **D**. The same strains were infected with ф37 (MOI=0.01) and cultures were passaged twice in LB medium (*n*=3). For each passage, bacterial (left) and phage (right) counts are shown. Constitutive modifications of the O-antigen prevented the replication of ф37 and provoked its extinction. **E-F**. Bacterial counts **(E)** and phage titers **(F)** in long-term experimental evolution *in vitro* revealed the stable coexistence of *S*.Tm with ф37 for at least 20 passages, *i.e.*, 200 bacterial generations, in LB (*n*=6). **G**. Detailed plaque assay results from the experiments presented in Figure 1 G-H and summarized in Figure 1I. The susceptibility of *S*.Tm to ф37was tested on 8 colonies from each biological replicate after the third passage; Yellow = clear plaques, susceptible clone; Orange = turbid plaques, heterogeneous phenotypic resistance linked to the expression of *gtrABC*; Red = no plaque, resistance linked to mutations in *btuB*. **H**. Control plaque assays in 24-well plates with ф37 and *S*.Tm reporter strains (genotypes indicated for each strain). ф37 formed turbid plaques on *gtrABC* ON cells. *S*.Tm WT carrying the *gtrABC*-*lacZ* transcriptional fusion was plated on LB agar X-Gal. Forty-eight blue colonies (*gtrABC* ON, black arrow) and 48 white colonies (*gtrABC* OFF, white arrow) were challenged with ф37. All the plaque assays performed with *gtrABC* ON colonies displayed turbid plaques, while the plaque assays performed with *gtrABC* OFF colonies displayed clear plaques. Six representative examples are depicted.

**Figure S2.**
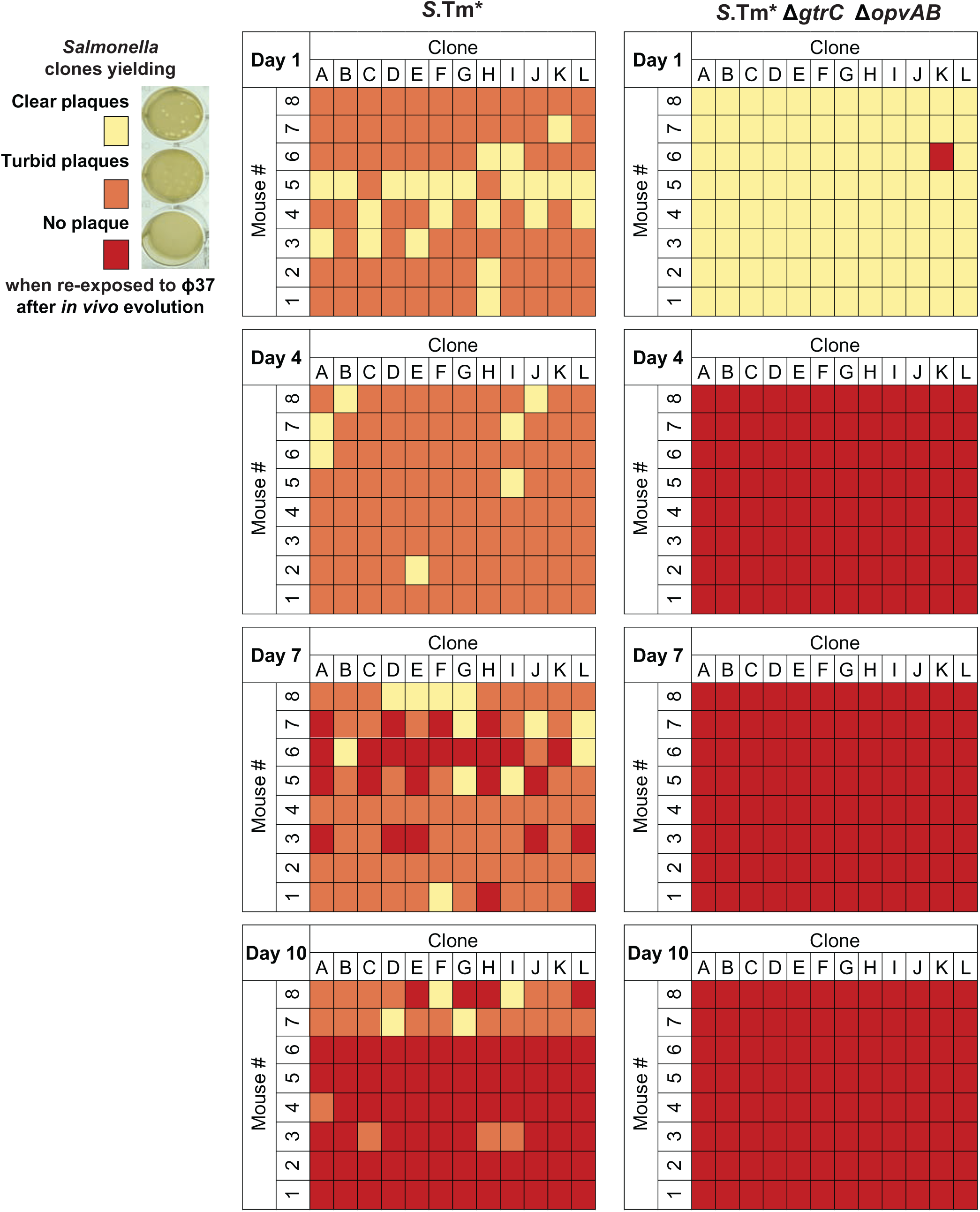
Plaque assays on isolated clones of *Salmonella* reveal that phase variation delays the fixation of *btuB* mutants. Detailed data summarized in Figure 2D. Individual clones from mouse fecal samples were tested for their susceptibility to ф37. Yellow = clear plaques, fully susceptible clone; orange = turbid plaques, partially resistant clone (GtrABC-modified O-antigen); red = no plaque, resistant clone (*btuB* mutants).

**Figure S3.**
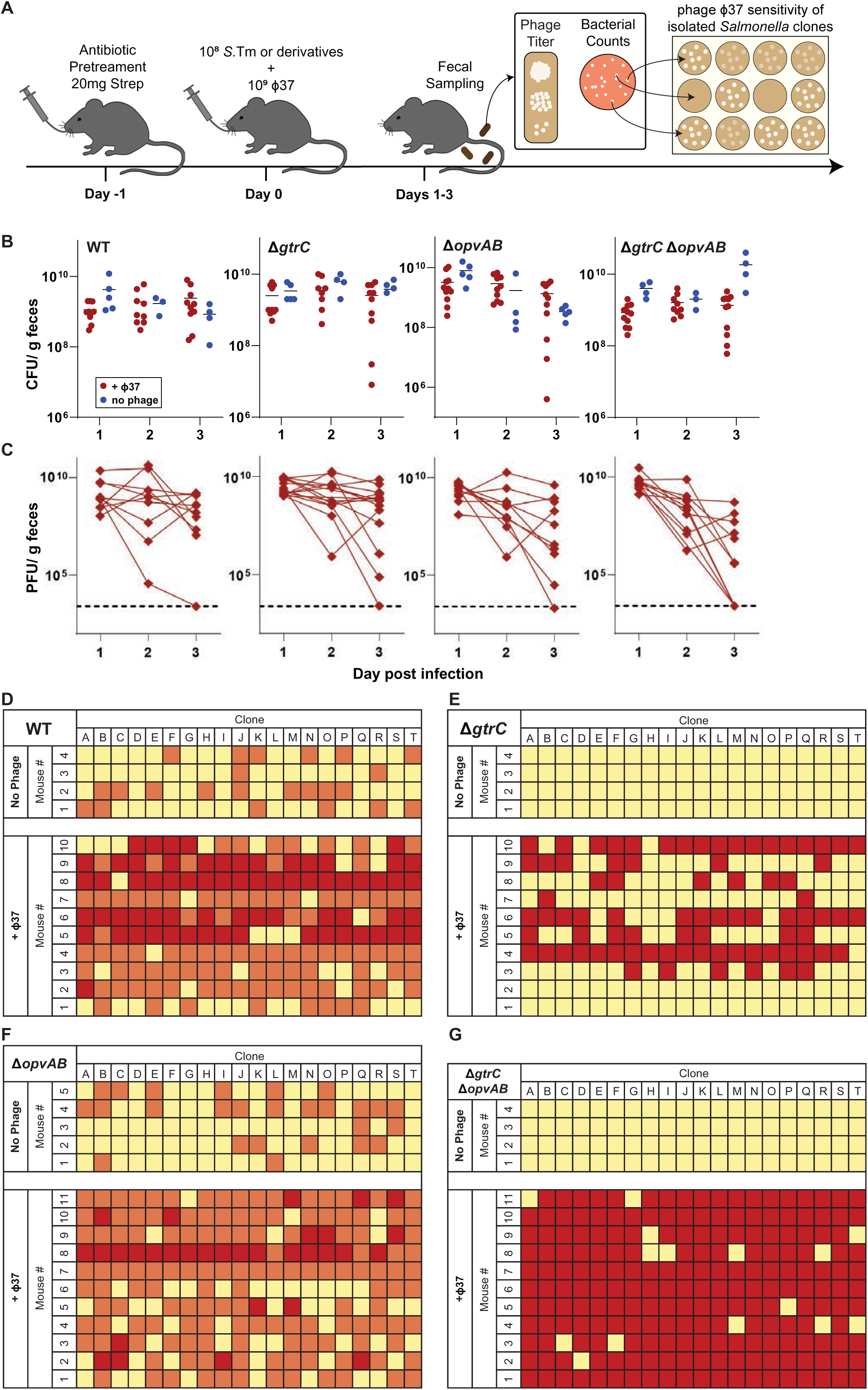
O-antigen phase variation by *gtrABC* and *opvAB* delays the fixation of genetic phage resistance in fully virulent *S*.Tm derivatives. A. Experimental setup. Streptomycin pretreated C57BL/6 Specific Pathogen Free (SPF) mice were infected with *S*.Tm WT, the mutants Δ*gtrC* or Δ*opvAB*, or the double mutant Δ*gtrC* Δ*opvAB* (10^8^ CFUs) and then received 10^9^ PFU of ф37 or exhausted LB as control. Bacterial **(B)** and phage **(C)** loads were quantified in fecal samples for 3 days. **D-E**, phage susceptibility of individual clones from each mouse was determined after 3 days. Yellow = clear plaques, fully susceptible clone; orange = turbid plaques, partially resistant clone (GtrABC-modified O-antigen); red = no plaque, resistant clone (*btuB* mutants).

**Figure S4.**
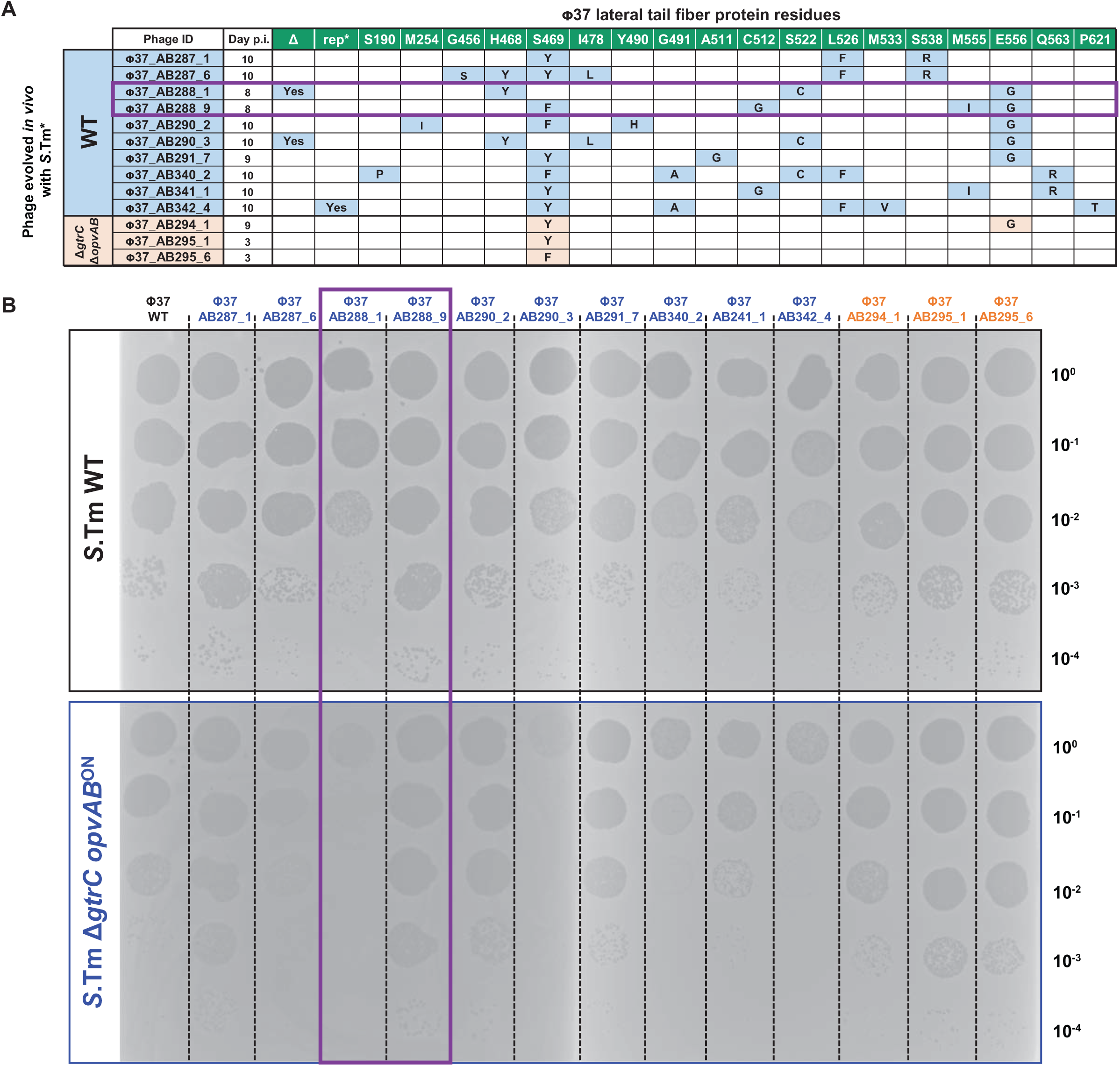
Infectivity of different *ltf* variants of ф37 on *opvAB*^ON^ cells. **A**. Table showing the *ltf* alleles of evolved phages isolated from mouse fecal samples (see Figure 2) at various days post infection (p.i.). **B**. Lysate of ancestral ф37 and *in vivo* evolved ф37 mutants were spotted on lawns of *S.*Tm WT and *S.*Tm Δ*gtrC opvAB*^ON^. Phages used in experiments presented in Figure 4 are highlighted in purple.

**Figure S5.**
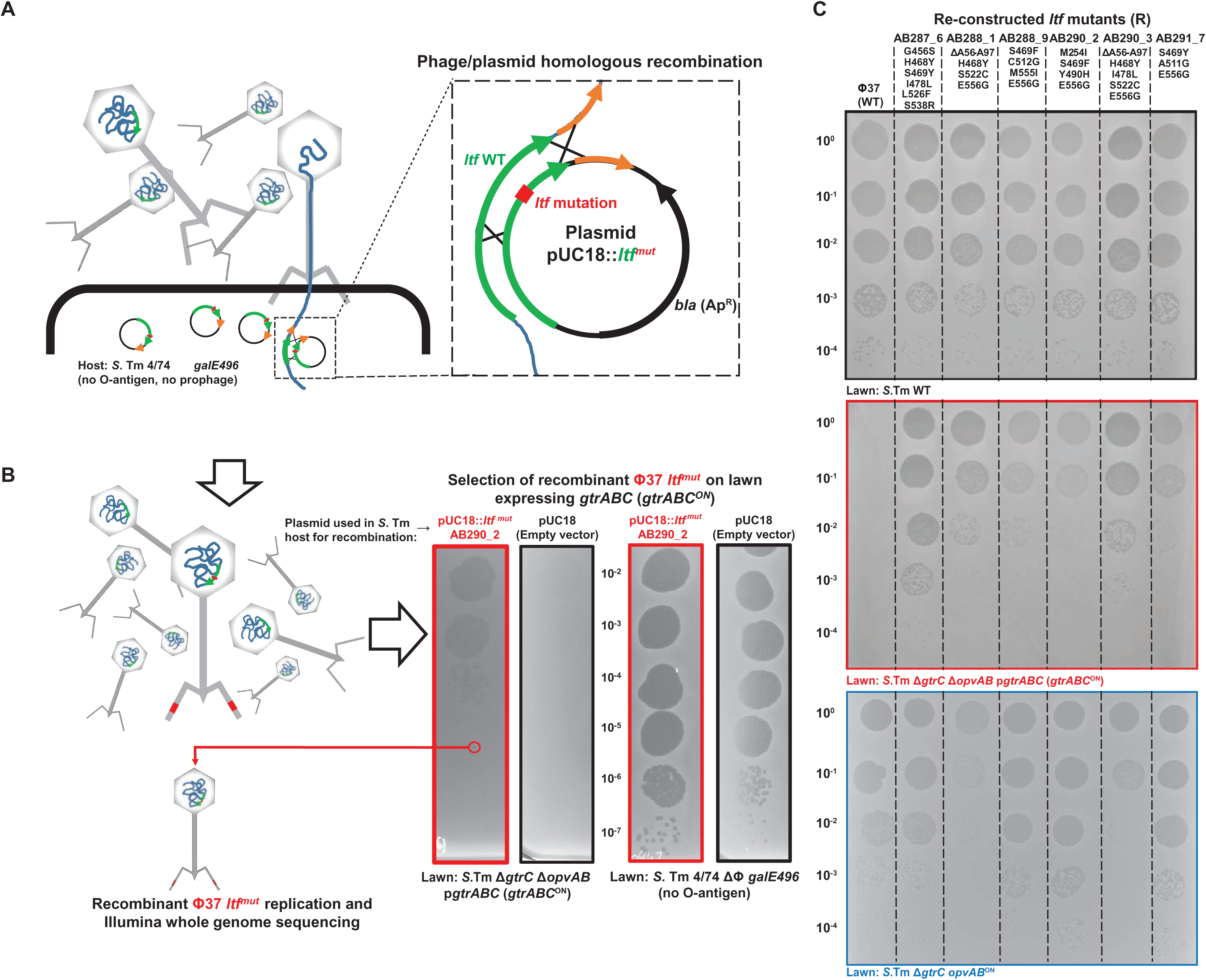
Re-construction and infectivity of *ltf* variants of ф37. **A-B**. Strategy for the re-construction of ф37 *ltf* variants. **A**. The *ltf* locus of a subset of the *in vivo*-evolved ф37 was cloned into plasmid pUC18. The resulting plasmids pUC18::*ltf*^mut^ were introduced into the prophage-free and O-antigen negative strain *S*.Tm 4/74 Δф *galE496*. The strains bearing the pUC18::*ltf*^mut^ plasmids or the empty pUC18 vector (negative control) were infected with the ancestral ф37 (MOI=5) to allow homologous recombination. **B**. The recombinant phages were selected by plaque assay on *S*.Tm *gtrABC*^ON^ (Δ*gtrC* Δ*opvAB* p*gtrABC*). Phages from individual plaques were amplified on the production strain 4/74 Δф *galE496*. Illumina whole genome sequencing confirmed the presences of the desired *ltf* mutations (**Dataset S2**). A representative example of re-construction (ф37_AB290_2) with plasmid pUC18::*ltf*^AB290_2^ is depicted. **C**. Assessment of the infectivity of the re-constructed ф37 *ltf* variants (R) on *S*.Tm, *gtrABC*^ON^ and *opvAB*^ON^ by plaque assay. Phage lysates were diluted and spotted on lawns of *S*.Tm WT, Δ*gtrC* Δ*opvAB* p*gtrABC* or Δ*gtrC opvAB*^ON^. The *ltf* mutations are indicated for each variant.

**Figure S6.**
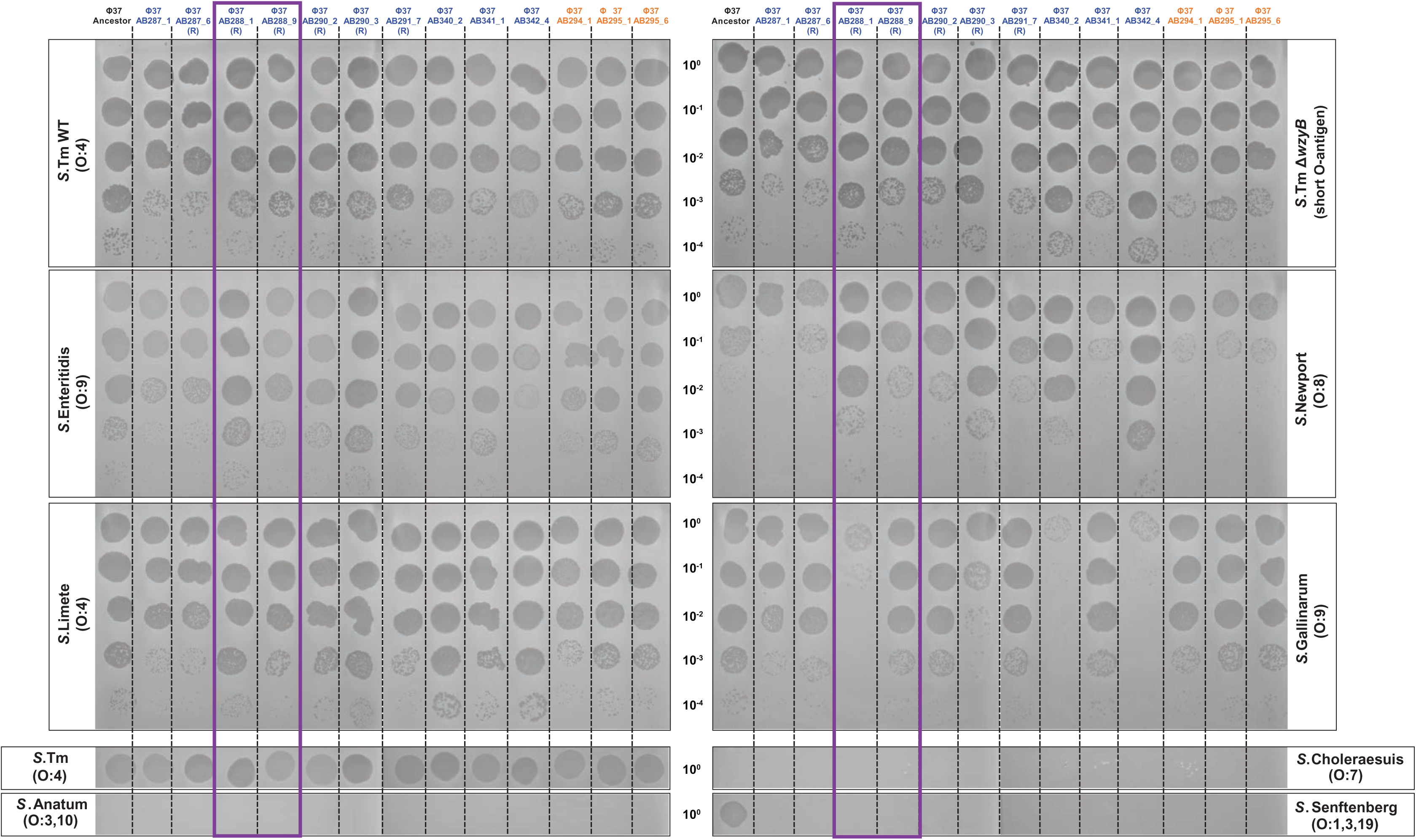
Infectivity of evolved phages and reconstructed phages on diverse *Salmonella* serovars. Lysates of the ancestral phage ф37, *in vivo-*evolved ф37 mutants and reconstructed *ltf* mutants (marked with (R) as presented in **Figure S5**) were spotted on lawns of *S.* Tm WT and different serovars of *Salmonella enterica*. Phages used in experiments presented in Figure 4 are highlighted in purple.

**Figure S7.**
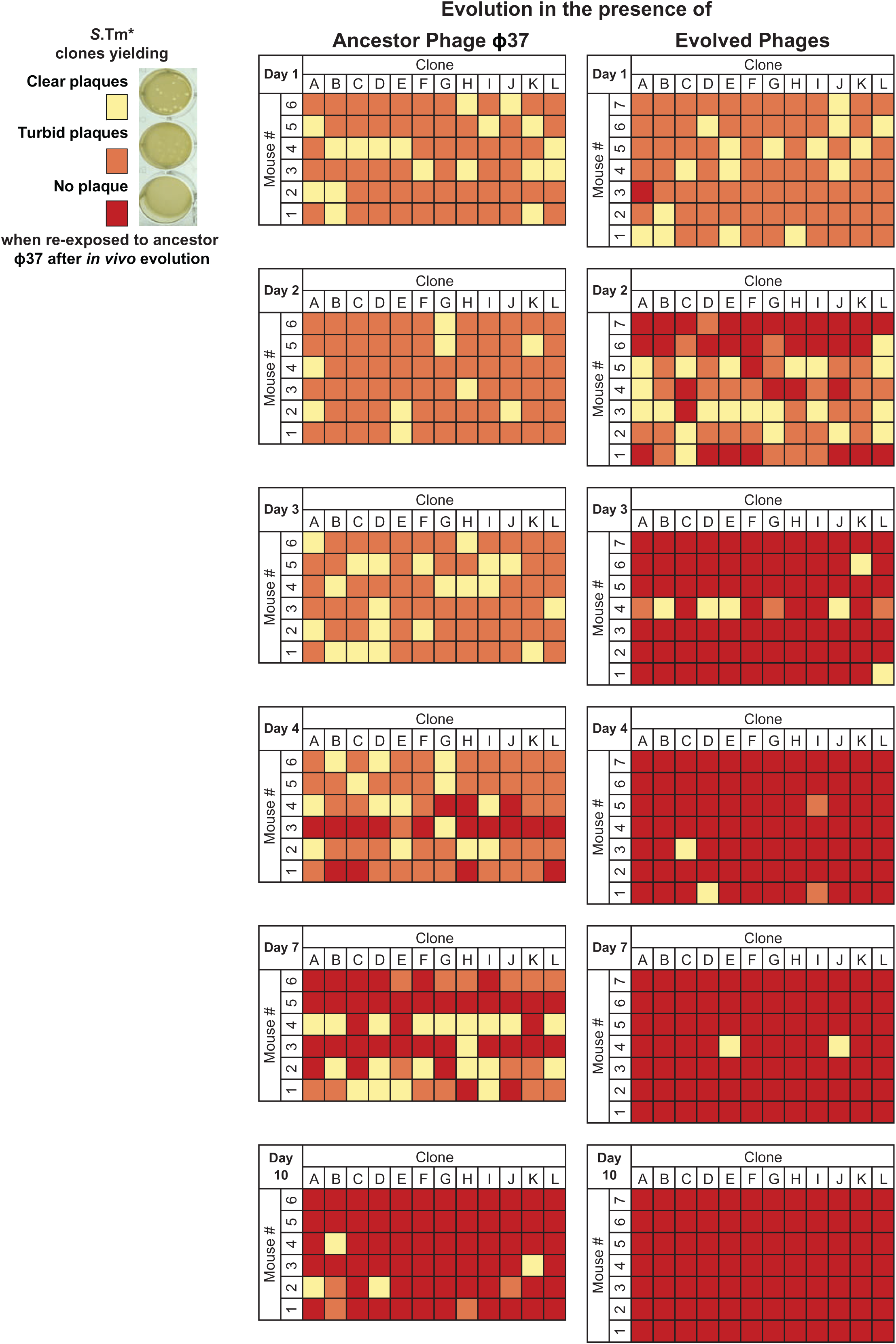
Plaque assays on lawns of isolated clones reveal the accelerated fixation of *btuB* mutants in the presence of evolved phages. Detailed data summarized in Figure 4D. Individual colonies from mouse fecal samples were tested for their susceptibility to ф37. Yellow = clear plaques, fully susceptible clone; orange = turbid plaques, partially resistant clone (GtrABC-modified O-antigen); red = no plaque, resistant clone (*btuB* mutants).

## Methods

### Culture conditions and transformation

All the resources used in this study are listed in **Table S1**. Unless stated otherwise, bacterial were grown from single colonies in autoclaved LB (10 g/L Tryptone, 5 g/L yeast extract, 10 g/L NaCl) at 37°C with shaking (200 rpm). Overnight (O/N) cultures are defined as 2 mL LB cultures grown at 37°C with shaking for 16-20 hours. Alternatively, strains were grown on solid LB agar plates (1.5% agar) or on MacConkey agar plates. When required, antibiotics were added to the media: streptomycin (Sm), 100 μg/mL; chloramphenicol (Cm), 6 μg/mL; kanamycin (Km), 50 μg/mL; ampicillin (Ap), 100 μg/mL. When required, inducers were added to the cultures: anhydrotetracycline (AHT) 0.5 μg/mL; L-arabinose 0.2% (w/v). For the growth of *E. coli* JKe201 **[38]**, 1,6-diaminopimelic acid (DAP, 0.1 mM) was added to the cultures. To visualize the *gtrABC* ON/OFF status of strains carrying the bistable *gtrABC-lacZ* transcriptional fusion **[24]**, bacteria were plated on LB agar plates supplemented with 100 μg/mL X-Gal and incubated O/N at 37°C.

For the preparation of electro-competent bacteria, strains were grown to exponential phase in salt-free LB (10 g/L Tryptone, 5 g/L yeast extract) and washed in cold distilled water. Bacterial transformation was performed by electroporation (2.5 kV) in 2 mm gap electroporation cuvettes with 0.1-3 µg of purified DNA, using the Gene Pulser Xcell eletroporator (BioRad).

### Cloning procedures

The construction of all the bacterial strains and their usage in each Figure is detailed in **Table S1**. Bacterial genomic DNA was isolated using the Quick-DNA Miniprep Plus Kit (Zymo). Bacteriophage DNA was isolated using the Phage DNA Isolation Kit (Norgen) and plasmids were isolated with the GenElute^TM^ Plasmid Miniprep Kit (Sigma). DNA purification from agarose gel or enzymatic reactions was performed with the NucleoSpin Gel and PCR clean up kit (Macherey-Nagel). PCRs were performed with Phusion DNA polymerase (Thermo Fisher Scientific), as specified by the manufacturer. When required DMSO (3%) and betaine (1 M) were added to the PCR reactions. Colony PCRs were performed with GoTaq® G2 Mastermix (Promega). Restriction enzymes and T4 DNA ligase were provided by New England Biolabs and Thermo Fisher Scientific.

### Bacterial genome editing techniques

The construction of all the bacterial strains is detailed in **Table S1**. All the *Salmonella* mutants were constructed in the fully virulent *Salmonella enterica* serovar Typhimurium SL1344 genetic background **[39]**. The wild type (WT) SL1344 from our collection (strain SB300), has been recently re-sequenced and compared with the reference genome **[40]**. Other *Salmonella* serovars, including strains from the SARB collection **[41, 42]**, were kindly provided by Prof. E. Slack (ETH, Zürich) and have been sequenced.

Gene deletions were performed with PCR fragments by D *red* recombination using the helper plasmid pKD46 and the template plasmids pKD3 and pKD4 **[43]**. To insert *gfp* downstream of the *opvAB* operon (strain *opvAB-gfp*) by D *red* recombination, plasmid pNAW52 **[44]** was used as donor for the *gfp-frt-aphT-frt* cassette. The P22 HT *105/1 int-201* transducing phage was used to transduce antibiotic-marked mutations in *S*.Tm **[45]**. When required, antibiotic resistance cassettes were removed with the FLP recombinase expressing plasmid pCP20 **[43]**.

To transfer the *opvAB^ON^* mutations (four GATC→CATG mutations in the *opvAB* promoter) into *S.*Tm SL1344 derivatives, the *opvAB* promoter from *S.*Tm 14028 *opvAB*^ON^ (SV6401) **[23]** was PCR-amplified and cloned into the allele replacement plasmid pFOK **[46]**. The resulting plasmid pFOK-*opvAB*^ON^, was mobilized from *E. coli* JKe201 into SL1344 by conjugation and merodiploid resolution was performed, as previously described **[46]**.

All mutations were confirmed by PCR and Sanger Sequencing (Microsynth AG, Balgach, Switzerland). Whole genome sequencing (WGS) by Illumina and/or Nanopore MinION was used to control several key strains, as specified in the bacterial strain list (**Table S1**).

### Phage manipulation and plaques assays

All the phages used are derivatives of the T5-like *Salmonella* phage ф37 (subfamily *Markadamsvirinae*, genus *Epseptimavirus*), listed in **Table S1 [14]**. Lysates of *Salmonella* phage ф37 and of its variants were prepared on the prophage-free strain *S*.Tm 4/74 Δф **[47]** carrying the defective *galE496* allele, linked to an *aphT* (Km^R^) cassette **[48]** (Strain MDBZ0942). MDBZ0942 does not produce O-antigen, and was used as host to prevent lateral tail fiber (Ltf) protein evolution during the passaging and lysate preparation of *in vivo*-evolved ф37 variants.

To obtain high-titer phage lysates, 10^4^ to 5.10^4^ Plaque Forming Units (PFUs) were mixed with 100 µL of MDBZ0942 O/N culture in 10 mL of warm (50°C) Top Agar (LB + 0.5% agar). The mixture was poured on a 12 cm x 12 cm square LB agar plate. After solidification the plate was incubated at 37°C O/N. The top agar layer was removed and suspended in 10 mL LB. After vigorous vortexing, the suspension was spun down (20 min, 4000*g*) and the supernatant was filtered (0.22 µm). All phage lysates were stored a 4°C with 1% chloroform.

Phage enumeration was performed using double agar overlay plaque assay **[49]**. For 8.6 cm diameter petri dishes, 5 mL Top Agar was used and for 12 cm x 12 cm square plates 10 mL was used. The reporter strain O/N cultures were diluted 1:50 in the warm Top Agar, which was poured onto a LB 1.5% bottom agar plate. After Top Agar solidification, 10 μL or 5 μL of diluted phage samples (10^0^ to 10^-8^ dilutions in LB or PBS) were spotted on the Top Agar surface. After drying, the plaque assay plates were incubated O/N at 37°C and phage titer (PFU/mL) in the original sample was deduced by counting distinct plaques from the appropriate dilutions.

For all the plaque assays presented in the figures, the phage lysates were adjusted to ∼10^7^ PFU/mL, serially diluted to 10^-4^ and 10 μL of dilutions 10^-4^,10^-3^,10^-2^, 10^-1^ and 10^0^ were spotted on lawns of the indicated reporter strain.

To screen numerous *S*.Tm clones for phage ф37 sensitivity, plaque assays were performed in a 24-well microplate format. Isolated *S*. Tm colonies on agar plates were picked and inoculated into the wells of a 96-well microplate, containing 50 µL LB. The bacteria were grown in microplates for 2 hours at 37°C with shaking at 200 rpm. The 50 µL cultures were mixed with 250 µL of warm Top Agar and poured in the wells of 24-wells microplates, containing already 500 µL of bottom LB agar 1.5%. After solidification, 20-50 PFUs of phage ф37 were applied on the Top Agar surface. After drying, plates were incubated O/N at 37°C and plaques presence/absence and morphology was recorded.

### Phage replication assay

20 μL of each bacterial O/N culture were diluted in 2 mL of LB medium in a 15 mL Falcon tube and incubated at 37°C with shaking. When the plasmid p*gtrABC* was used, Ap was added to the cultures. After 2h incubation (exponential phase), 200 μL of each culture were transferred into 1.5 mL tubes. About 10^3^ PFU/mL (MOI ∼10^-5^) of ф37 (or its derivatives) were added to each tube. After 6 hour incubation (37°C, 200 rpm), replication was stopped by adding 20 μL of chloroform. After vortexing and centrifugation (3 min, 14000 rpm) the phage titers were determined by plaque assay on lawns of *S*.Tm Δ*wzyB*. The phage replication fold was calculated by dividing the final phage titer by the input phage titer added at T_0_. Each experiment was performed at least twice with biological triplicates.

### Phage adsorption assay

5 mL of LB were inoculated with 50 μL of *S*.Tm O/N culture and incubated at 37°C with shaking for 2h 40 min. Each culture and a control (5 mL LB without bacteria) were inoculated with 5 * 10^6^ PFUs of ф37 (MOI 0.01-0.001) and incubated for another 15 min. 200 μL of culture were collected, spun down and treated with chloroform. Plaque assay was performed to determine the free phage titer and adsorption efficiency was calculated by dividing the final free phage titer by the input phage titer added at T^0^. Each experiment was performed at least twice with biological triplicates.

### Growth curves in the presence of phages

500 μL of LB were inoculated with 2.5 μL of O/N *S*.Tm cultures and infected with 10^5^ PFU/mL of phage ф37 (final MOI = 0.01). 200 μL were transferred in a 96-well microplate. The plate was incubated overnight at 37°C with shaking in a Synergy H4 plate reader (BioTek), measuring Optical Density at 600 nm (OD_600_) every 10 minutes. The data was then corrected by subtracting the background (LB) OD_600_ before plotting. Each experiment was performed with biological triplicates.

### Experimental evolution in LB

5 mL of LB in 50 mL tubes were inoculated with the corresponding strain (10^6^ CFU/mL) and phage (10^4^ PFU/mL). After 24h of growth at 37°C and with shaking, the culture was diluted a thousand times in a new tube containing 5 mL of LB. For each passage, CFUs were determined by plating dilutions of each replicate on LB plates. For phage titer determination, 1 mL of culture was filtered (0.22 μm) and PFU/mL was determined by plaque assay, using the reporter strain *S*.Tm WT. Each experiment was performed twice with biological triplicates and appropriate controls (no phage).

### Re-construction of *ltf* variant phages

The *ltf* alleles found in *in vivo*-evolved phages were transferred into the ancestral ф37 using the technique developed by Ramirez-Chamorro and colleagues **[30]** (**Figure S5A&B**). DNA fragments carrying the *ltf* mutations were PCR-amplified from ф37 variants AB287_6, AB288_1, AB288_9, AB290_2, AB290_3 and AB291_7. The resulting amplicons were cloned into pUC18 and the resulting plasmids (pUC18-*ltf*^Mut^) were introduced into strain MDBZ0942. O/N cultures of each transformed strain carrying the pUC18-*ltf*^Mut^ or the empty pUC18 (negative control) were diluted 10 times in LB. After 30 min incubation at 37°C with shaking, the cultures were infected with ф37 WT (MOI=5), and incubation was continued for 20 hours. Culture supernatants were filtered (0.22 μm) and serial-diluted to 10^-8^. To select for the recombinant *ltf*^Mut^ phages, the dilutions were spotted on top agar lawns of *S*.Tm expressing constitutively *gtrABC* (strain *gtrABC*^ON^, MDBZ1561). Single isolated plaques obtained on the *gtrABC*^ON^ lawns were picked and passaged twice on strain MDBZ0942 (O-antigen-free strain). PCR and Sanger sequencing with primers OMD22_400 and OMD23_001 confirmed the presence of the desired *ltf* mutations. Finally, Illumina WGS confirmed the *ltf* mutations and the absence of accessory mutations in the re-constructed ф37 *ltf*^Mut^.

### O-antigen analysis by SDS-PAGE and silver staining

200 μL of O/N cultures were spun down and bacteria were suspended in 500 μL of Laemmli loading buffer 1 X (10 mM Tris-HCl, 2 % SDS, 3 % DTT, 10 % glycerol, 0.1% Bromophenol Blue, pH 6.8). 10 μL of Proteinase K (20 mg/mL) were added and the suspension was incubated for 3 h at 58°C. The lysate was boiled for 5 min and spun down. 10 μL were loaded on a 13% polyacrylamide (PAA) Tricine-SDS separating gel topped with 4 % PAA stacking gel **[50]**. After electrophoresis (45 min, 200 V), the gel was washed twice for 5 min with water and soaked in Fixer Solution (30 % ethanol, 10 % acetic acid) O/N. The gel was then incubated 5 min in 25 mL of oxidizing solution (Fixer solution + 0.35% periodic acid) and washed twice 5 min with water. LPS bands were revealed by silver staining using the Pierce^TM^ Silver Stain Kit, according to the manufacturer’s instructions (Invitrogen).

### Immunostaining and flow cytometry

All flow cytometry experiment were conducted at the Biozentrum FACS Core facility on a BD LSRFortessa Cell Analyzer. For *S*.Tm O-antigen (O:12) immunostaining, 1 mL of O/N LB culture was spun down and bacteria were re-suspended in 1 mL BSA (1% Bovine Serum Albumin in PBS) and incubated on ice for 1h. The cells were spun down again, re-suspended in 1 mL PBS-BSA and 1 μL of STA5 Anti-O:12 antibody (human recombinant monoclonal IgG2 anti-O:12-0; 6 µg/mL) was added **[51]**. After 1h incubation on ice, the bacteria were washed twice with 1 mL PBS-BSA and then re-suspended in 1 mL PBS-BSA. 1 μL Alexa Fluor-647 Goat Anti-Human IgG antibody was added. After 1h incubation on ice, bacteria were washed with 1 mL PBS-BSA and then re-suspended in 1 mL PBS-BSA.

The bacteria suspensions were diluted 1:100 in filtered PBS for flow cytometry analysis. The analysis of the *gfp* expressing reporter strains was conducted without immunostaining. Flow cytometry data were analyzed using FlowJo (BD).

### Ethics

All animal experiments were approved by the legal authorities (Basel-Stadt Kantonales Veterinäramt, licences #30480 and #33580) and follow the 3R guidelines to reduce animal use and suffering to its minimum.

### Murine infection models

C57BL/6 mice used in this study were either conventional Specific Pathogen Free (SPF), harboring a complex microbiota, or LCM (Low Complexity Microbiota) harboring a simplified microbiota generating a reduced colonization resistance against *S*.Tm **[26]**. All mice were housed and bred at the Werk Rosental Animal Facility of the University of Basel.

Eight- to twelve-week old (males and females) SPF and LCM mice were pre-treated with 25 mg streptomycin 24h before *S*.Tm infection *per os*. Microbiota depletion by antibiotic pre-treatment allowed a robust and stable colonization with the Sm^R^ virulent strain SL1344 and its derivatives **[28]**. For short-term (3 days) infection experiments with fully virulent *S*.Tm, streptomycin pre-treated SPF mice were used. For long-term infection model (10 days), pre-treated LCM mice were infected with the attenuated *S.*Tm Δ*ssaV* mutants (*S*.Tm*)**[25]**.

Bacteria and phages were inoculated into mice *per os*. *Salmonella* inocula (10^8^ CFUs in 50 µL) were prepared as follows: *S*.Tm strains were grown O/N in 2 mL of LB with the appropriate antibiotics, diluted 1:20 and grown again in 2 mL LB for 4h at 37°C with shaking. Cells were washed twice with PBS and re-suspended in PBS. Phage inocula (10^9^ PFUs in 100 µL) were given to the mice 30 min after *S.* Tm infection. Phages were prepared in LB as described above on the O-antigen-free strain MDBZ0942. For “No phage” mock treatments, 100 µL of phage-free exhausted LB were used. Exhausted LB was prepared by filtrating (0.22 µm) an early stationary culture of MDBZ0942 in LB.

### Bacteria and phage enumeration from feces

Feces were collected on a daily basis. Droppings were weighted (5-80 mg) and homogenized in 1 mL PBS with 2 glass beads at 25 Hz for 3 min in a Qiagen Tissue Lyser II. The suspension was serial-diluted to 10^-6^ and 10 µL of each dilution were spotted on McConkey agar plates supplemented with the appropriate antibiotics. After O/N incubation at 37°C isolated colonies were counted and bacterial load was defined as Colony Forming Units *per* gram of feces (CFU/g Feces).

For phage enumeration, 250 µL of homogenized feces were vortexed for 15 sec with 20 µL chloroform. After centrifugation (3 min, 14000 rpm), the supernatant was serial diluted in PBS and phage enumeration was carried out by plaque assay on a phage susceptible reporter *S*.Tm strain. After O/N incubation at 37°C, isolated plaques were counted and viral loads were defined as Plaque Forming Units *per* gram of feces (PFU/g Feces).

### Genome sequencing, assembly and annotation

Short-read Illumina sequencing was carried out by SeqCenter (Pittsburgh, USA). Oxford Nanopore MinION long-read sequencing was performed in house, using the MinION Flow Cells R10 (reference 10.4.1) and Rapid Barcoding Kit 24 V14 (Oxford Nanopore). Base calling was performed using Guppy v6.5.7+ca6d6af, and the base calling model was dna_r10.4.1_e8.2_400bps_sup (Oxford Nanopore).

Bacterial and phage genomes were assembled using Unicycler v0.5.0 (for short or combined short/long reads) **[52]** or with Flye v2.9.1 (only for long reads) **[53]**, with the default settings. Genomes were annotated using Prokka v1.14.6 **[54]**.

For variant-calling of the *de novo* sequenced strains, reads were aligned on the SL1344 reference genome (Genbank: NC_016810) with BWA-MEM v0.7.17.2 **[55]**. SNPs and Indels were identified using LoFreq v2.1.5 **[56]**.

The genome of bacteriophage ф37 **[40]** and its variants were visualized and aligned with SnapGene v4.0.3 and all mutations are reported in Supplementary dataset S2. All the raw sequencing data and assemblies will be made available on the NCBI Bio project database.

### Statistical analysis

Plots and statistical analyses were generated with Microsoft Excel 2016 and GraphPad Prism 9.3.1.471.

## Acknowledgements

We would like to acknowledge the labs of Jay Hinton (University of Liverpool, UK), Marjan Van der Woude (University of York, UK), Wolf-Dietrich Hardt and Emma Slack (ETH Zurich, Switzerland), and late Josep Casadesús for their kind contribution with strains and reagents.

We would like to thank Claudia Igler (University of Manchester) and the members of the Diard, Perez and Neher groups involved in the discussion of results and the sharing of ideas as well as the teams at the core facilities of the University of Basel for their valuable technical support.

## Fundings

NW, AB, AR and NAB were supported by SNSF professorships # PP00P3_176954 and PP00P3_213978 allocated to MD.

NW was also supported by a Microbial grant from the Gebert Rüf Stiftung # GRS–093/20 allocated to MD.

LL was supported by a multi-investigator grant from the BRCCH partially allocated to MD.

LR was supported by the ERC Consolidator grant ECOSTRAT #101002643 allocated to MD.

AH was supported by SNSF Ambizione #PZ00P3_180085.

VD was supported by the SNSF grant 310030_188547.

## Author contributions

NW: conceptualization, methodology, experiments, writing, supervision, review and editing.

AB: methodology, experiments, writing, review and editing.

LL, AR, NAB, CS, LR, VD: methodology, experiments, review and editing.

AH: methodology, review and editing.

MD: conceptualization, methodology, writing, review and editing, project administration, supervision and funding acquisition.

## Notes

### Competing Interest Statement

The authors have declared no competing interest.

